# Automated assembly of protein complexes from cryo-EM maps with structure-informed Monte Carlo Tree Search

**DOI:** 10.64898/2026.05.16.725663

**Authors:** Rohit Dilip, Songrong Jeff Qu, Zhen Chen, David Van Valen

## Abstract

Structural cell biology aims to visualize functional molecules as they carry out their biological roles in their native cellular context. However, macromolecular complexes *in situ* have thus far been resolved predominantly at intermediate resolutions, complicating protein identification and structural modeling due to the vast combinatorial space of possible components within a proteome. Here, we developed Cryosearch, a GPU-accelerated framework for automated assembly of macromolecular complexes from proteome-scale monomer libraries. Cryosearch implements Monte Carlo tree search with correlation-based rewards to identify combinations of protein domains that collectively best explain density maps. This approach enables autonomous *de novo* assembly of molecular complexes from intermediate-resolution maps.

## INTRODUCTION

Understanding biological processes often requires identifying the proteins involved and visualizing how they carry out their functions in their native environment, ideally at high resolution^1,2^. Advances in cryo-electron microscopy (cryo-EM) have enabled the determination of protein structures while preserving biological samples in hydrated states^3^. Recently, cryo-electron tomography (cryo-ET) combined with subtomogram averaging has emerged as a promising approach for resolving macromolecules *in situ*^4,5^. However, structures at near-atomic resolution (<4 Å) has largely been restricted to large macromolecular complexes, such as ribosomes, or repetitive cellular structures^6–8^. Due to limited throughput and intrinsic molecular heterogeneity, reconstructions of complexes within intact cells remain at intermediate resolutions (4–10 Å), where interpreting density maps and identifying protein components are challenging because the number and identity of constituent proteins are typically unknown and must be inferred from the entire proteome^9,10^. Addressing this challenge is essential for achieving visual proteomics—the ability to visualize and identify proteins from the proteome directly within cells^1,2^.

Current manual modeling approaches often require expert-guided segmentation of the density map into putative domains, followed by monomer assignment based on recognizable tertiary folds or by direct model building when side chains are resolved^11–14^. However, the peptide backbones could only be traced comprehensively at sub-4.5 Å resolution while side chains typically become discernible only at sub-4 Å resolution^9^, making intermediate-resolution maps difficult to model. Even in cases with sufficient resolution, this divide-and-conquer strategy is challenging to scale, as it requires iterative trial and error, substantial human intervention, and is inherently non-systematic^9,15^.

Correlation-based docking using proteome-wide databases predicted by AlphaFold offers an unbiased approach to automate the modeling and identification process at intermediate resolution (4-10 Å) (Fig. 1a)^10,16,17^. However, docking large numbers of candidate models into complex maps yields many false positives (Extended Data Fig. 1), requiring extensive manual evaluation of individual candidates^10^ and their combinations to explain the entire map. Currently, the field lacks methods to autonomously sample this vast combinatorial space at the proteome scale while efficiently converging on plausible solutions. Here, we present Cryosearch, a GPU-accelerated method for autonomous identification and modeling of proteins in large macromolecular complexes across the proteome. By combining correlation-based docking with Monte Carlo tree search (MCTS), Cryosearch efficiently samples combinations of protein domains that collectively best explain cryo-EM density maps.

**Figure 1.**
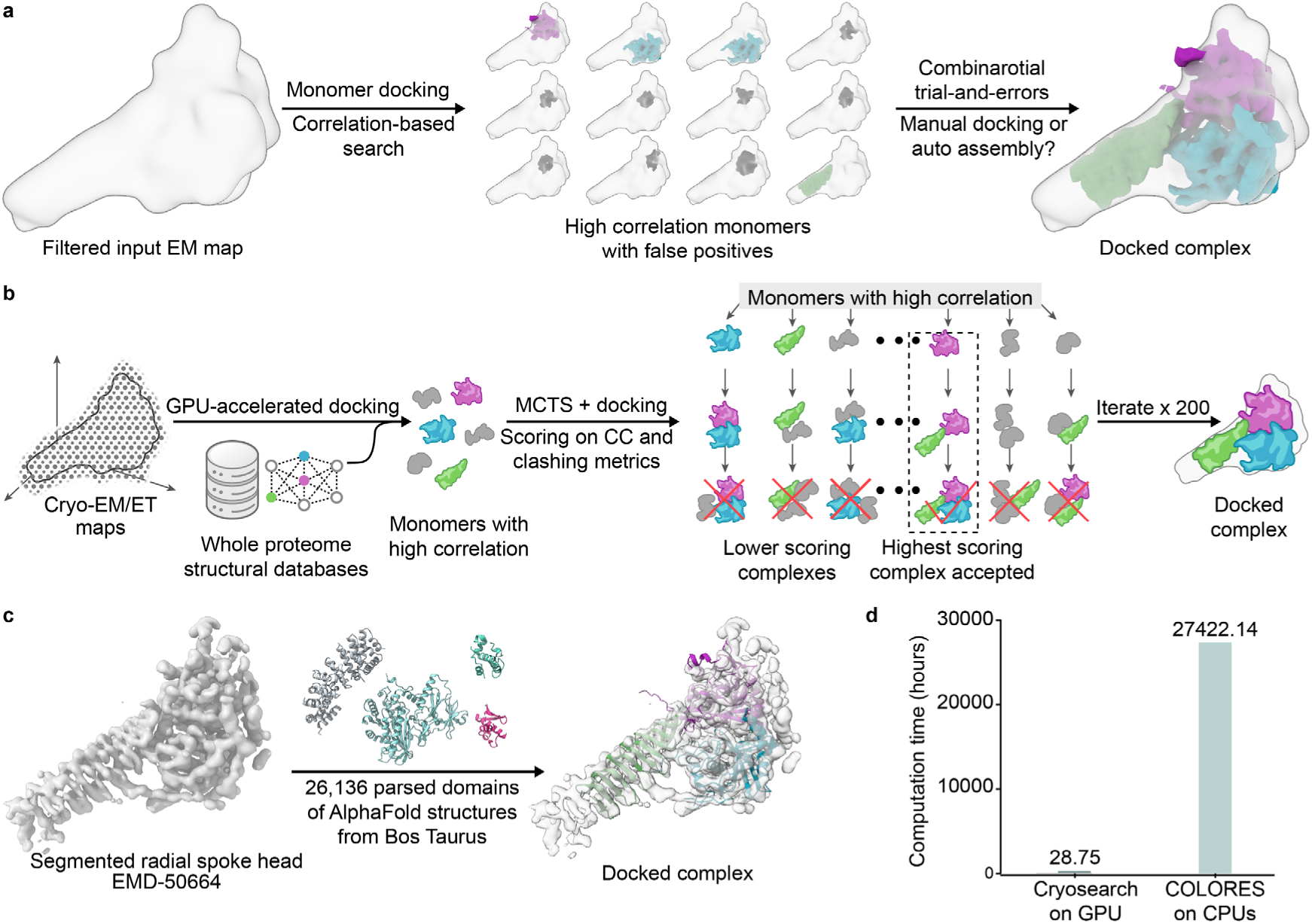
Cryosearch enabled proteome-scale identification and autonomous assembly of cryo-EM maps. **(a)** A schematic illustrating a conventional workflow of modeling multi-component complexes from density maps. The main challenges include false positives from monomer docking and a large combinatorial search space. **(b)** Overview of Cryosearch, which combines GPU-accelerated docking and Monte Carlo tree search (MCTS) for multimer assembly. **(c)** Application of Cryosearch to the segmented radial spoke head (EMD-50664) using 26,136 parsed domains from the Bos taurus AlphaFold proteome. **(d)** Comparison of computation time for proteome-scale monomer docking by the GPU-accelerated Cryosearch (NVIDIA RTX A6000) and the CPU-based COLORES (Intel Xeon E5-2687W v3 CPUs).

**Figure 2.**
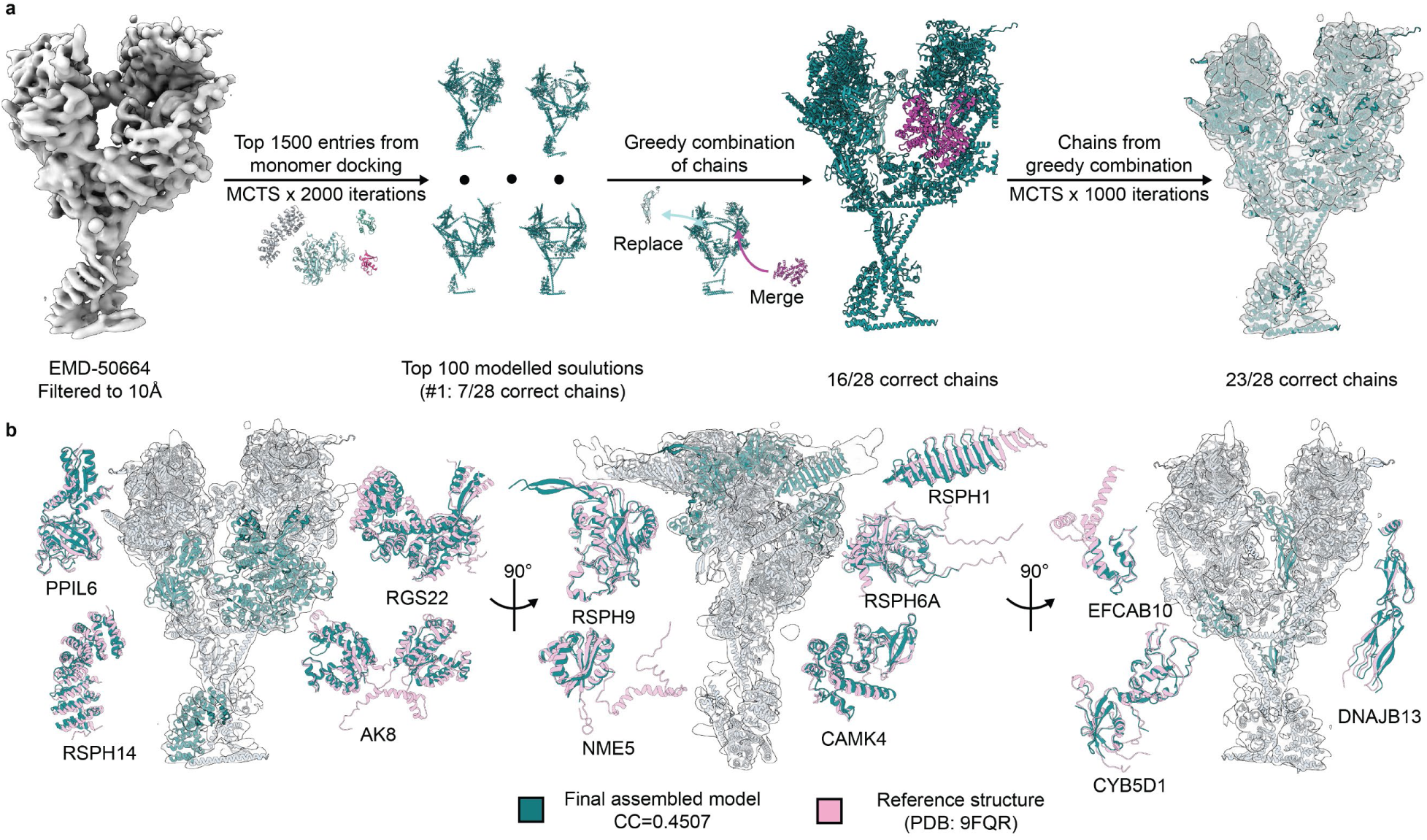
Cryosearch enabled autonomous modeling of large protein complexes with 20+ components. **(a)** A workflow for large-complex modeling of the radial spoke 1 head (EMD-50664), including initial MCTS assembly of partial solutions, greedy combination of chains from top assemblies, and a final MCTS refinement. 23 out of 28 correct chains were assembled autonomously (see discussion on the remaining components in the text and Fig. S7). **(b)** Alignment of the final assembly with the reference model (PDB: 9FQR) and correctly identified components.

## METHODS

We employed Monte Carlo tree search (MCTS), a decision-making algorithm that guides search for possible combinations of protein models that collectively best explain an input map, analogous to its applications in board games, such as AlphaGo. Cryosearch has three key components. First, we implemented correlation-based docking of PDB models into density maps in Fourier space and parallelized translational and rotational searches on a single GPU (Fig. 1b). Second, we used the cross-correlation score as the reward function of MCTS to assemble multi-subunit complexes that maximize the agreement between the model and map, while penalizing steric clashes (Fig. 1b). We envisioned that false positives might transiently improve the cross-correlation score but ultimately block placement of correct models, and thereby be disfavored by MCTS because they lead to lower collective cross-correlation scores. Third, for symmetric complexes, we optionally restricted late-stage proposals using nearest-neighbor searches in an embedding space derived from a global protein structure tokenizer^25^, thereby reducing the number of candidate identities explored (Extended Data Fig. 9).

## RESULTS

We first tested the feasibility of our MCTS strategy using a segmented map of the radial spoke from bovine sperm flagella (EMD-50664) (Fig. 1c)^18^. To perform an unbiased proteome-wide search, we parsed all bovine protein models predicted by AlphaFold based on Predicted Aligned Error (PAE) scores (also > 50 amino acids) (See **METHODS**)^16,19^. The resulting parsed bovine library contained 26,136 domains, which we docked individually into the map. We observed that GPU acceleration substantially reduced computation time, with monomer docking on a single GPU requiring a runtime ∼10^3^-fold faster than that of the CPU-based docking software COLORES (Fig. 1d)^17,20^. We then applied Cryosearch to the top 30 domains to identify combinations that maximize the cumulative cross-correlation. Cryosearch recovered two trimeric assemblies composed of either RSPH1/RSPH4A/RSPH9 or RSPH1/RSPH6A/RSPH9, both of which clearly match the secondary and tertiary structural features in the map (Fig. 1c and Extended Data Fig. 2). Both solutions were plausible because RSPH4A and RSPH6A are highly homologous. Comparison with the ground-truth model comprising RSPH1/RSPH6A/RSPH9 showed strong agreement in both the positions and orientations of the docked model within the EM map (Extended Data Fig. 2). Together, these results indicated that Cryosearch can efficiently and autonomously identify combinations of domains to account for density maps of complexes.

To evaluate scalability and convergence, we assessed Cryosearch across different numbers of MCTS iterations (50-500), which determines the number of combinations sampled before the output is generated (Extended Data Fig. 3a). Ten independent runs were performed at each iteration setting, and the top 5 outputs from each run were examined for accuracy. At 200 iterations, the correct trimer was consistently identified as the best model with the highest correlation score across all 10 independent runs, corresponding to a 100% success rate (Extended Data Fig. 3a). However, only 17 of the 50 top models (top five per run) were correct, resulting in an overall recovery rate of 34%. Increasing the number of MCTS iterations to 500 improved the recovery rate to 96% (48 out of 50 models) while maintaining runtime below 5 hours per run (Extended Data Fig. 3a-b). These results indicate that Cryosearch is a robust and efficient approach to converging on accurate models of protein complexes with sufficient trials.

We next tested the limits of Cryosearch for modeling large protein complexes at intermediate resolution, addressing the general challenge of protein identification in cryo-ET maps. The map of the radial spoke 1 head (EMDB-50664), modeled by 15 protein entries and 28 chains in the PDB model (PDB: 9FQR), was filtered to 10 Å to obtain an intermediate-resolution reconstruction (Fig. 2a)^18^. We first docked the parsed bovine database of 26,136 domains into the full map and ranked the cross-correlation scores (Extended Data Fig. 4a). Next, we conservatively retained the top 1,500 monomer domains for MCTS (Fig. 2a), resulting in a substantially larger search space compared to the segmented map (Fig. 1c).

We used MCTS to assemble a 40-component complex, which overestimated the number of domains in the map since the true number was generally unknown prior to modeling. Using 2,000 MCTS iterations to limit runtime to 48 hours, the top-ranked model recovered 6 out of the 15 protein entries and 7 out of the 28 chains (Extended Data Fig. 5). However, we observed that different correct proteins and chains were recovered across multiple top-ranked models (Extended Data Fig. 5), suggesting that they may collectively better represent the correct assembly. We therefore developed a refinement procedure to combine these partial models. Starting with the best model, chains from the top 100 models were either added or used to replace existing chains at overlapping positions if such actions improved the overall cross-correlation. This greedy combination increased recovery to 12 of 15 protein entries and 16 of 28 chains (Extended Data Fig. 6). Extending the greedy combination to the top 500 models yielded similar recovery of protein entries (12/15) and chains (17/28) (Extended Data Fig. 6), suggesting that the optimization had converged. Analyses of the missing entries revealed several intrinsic limitations of intermediate-resolution modeling. We noted that ROPN1L was ranked 8037^th^ in monomer docking, likely due to the lower resolution at the periphery of the map (Extended Data Fig. 4a-b). RSPH3 and DYDC1 contain extended single α-helix structures (Extended Data Fig. 4c). Although our model docked other single α-helix domains with similar lengths (Extended Data Fig. 7b), we generally rejected them since unambiguous identification of such structures at 10 Å is not possible.

Finally, we performed a second MCTS focused on the domains identified during greedy combination (46 in total), using their poses as prior knowledge to guide the search toward improved combinations. The final output showed an overall good fit of 23/28 chains to the density map, with tertiary and secondary structural features well-matched, only missing the five chains from ROPN1L, RSPH3 and DYDC1 (Fig. 2b and Extended Data Fig. 7a). Our 40-chain models contain six ambiguous single α-helix chains in the regions of RSPH3 and DYDC1 (Extended Data Fig. 7b). Another six false positives were included and were ranked 31, 35–38, and 40 based on cross correlation (Extended Data Fig. 7c). The rank 35 chain corresponds to an RBM26 domain not present in the original reference model but matches well with the density (Extended Data Fig. 7c). The remaining five false positives either do not match secondary structures to the map or are single α-helix domains, allowing rejection upon manual inspection. Together, these results demonstrate that Cryosearch can autonomously and accurately model large protein complexes at intermediate resolution.

## DISCUSSION

Our results collectively demonstrated that Cryosearch enables proteome-scale modeling of protein complexes from intermediate-resolution cryo-EM maps by efficiently navigating the combinatorial search space using MCTS and GPU-accelerated docking. This framework could be further accelerated through parallelization across multiple GPUs in the future. Although MCTS does not converge on a single perfect model, it reliably captures multiple correct components across different trials (Extended Data Fig. 5), effectively narrowing the solution space from a vast pool of candidates. This efficiency arises because partial assemblies that improve cumulative cross-correlation are preferentially expanded during tree search. In this framework, correct components are consistently sampled across independent search paths because they locally improve the fit to specific regions of the map (Extended Data Fig. 5). In contrast, false-positive components tend to interfere with the placement of correct components by occupying incompatible regions, thereby reducing cumulative cross-correlation and disfavoring those paths during tree expansion. Once the candidate space was reduced to a manageable subset of the proteome and intermediate solutions were established (Extended Data Fig. 5), the greedy combination and the second MCTS efficiently refined the search and accurately modeled large complexes. Cryosearch, therefore, enables rapid and systematic generation of structural models for large complexes at intermediate resolution, a challenging task even for experts, by turning model building into a collaborative exercise between humans and algorithms.

AlphaFold-based approaches have greatly advanced structure prediction, but remain challenging to scale to large assemblies because accuracy decreases with increasing chain number and GPU memory requirements become limiting^21^. In addition, current automated or generative cryo-EM modeling methods often rely on protein sequence information for chain identification and tracing at intermediate resolution^14,22^. In contrast, Cryosearch enables *de novo* modeling from intermediate-resolution maps when side-chain information is unavailable and structural features are ambiguous.

The increasing amount of high-quality cryo-ET and cryo-EM data suggests a bright future for data-driven solutions to this problem. While our approach leverages a clear inductive bias for the problem – that monomer structures must be sampled from the proteome – additional data may open up the door to purely generative solutions. Cryosearch’s success also suggests that interleaving search with deep learning is a promising alternative. The improved search efficiency we observed when using protein tokenizers to bias proposals during MCTS for symmetric complexes is just one example (Extended Data Fig. 9). Similar to modern game solvers like AlphaGo^24^, additional data - both real and synthetic – could empower deep learning models to bias the search routine, reducing the search space from the full proteome to a smaller number of plausible monomers. With sufficient data, search algorithms could be trained via reinforcement learning to create neural reward functions that consider the full horizon over building a complex.

Several factors may limit Cryosearch’s performance. Although ∼10 Å resolution is generally sufficient to recognize protein folds (Fig. 2), local resolution can vary substantially, and peripheral regions of cryo-EM maps often suffer from lower resolution^22,23^. Such limitations may be mitigated by focused refinement. In addition, our approach relies on well-defined tertiary structures predicted by AlphaFold, and therefore excludes low-confidence regions, consistent with standard practices in expert-guided modeling^11^. Proteins dominated by single α-helices and homologous domains remain difficult to distinguish at intermediate resolution due to the absence of side-chain information, representing an inherent limitation shared with manual interpretation. These limitations, however, reflect fundamental constraints of the underlying data rather than shortcomings unique to Cryosearch. Overall, at ∼10 Å resolution, Cryosearch achieved modeling performance that remains challenging for expert-guided approaches, given the limited structural information and the large combinatorial space of macromolecular assemblies with more than 20 components.

## ACKNOWLEDGMENTS AND AUTHOR CONTRIBUTIONS

We thank Bricelyn Strauch for contributing illustrations. We thank Lior Pachter, Pietro Perona, Samuel Arnesen, Cory Acreman, Julian Braxton, Xiang Feng, Gohta Aihara, Tiffany Hung, Ada Ates, Sinan Ozbay, and William Arnesen for helpful discussions and feedback. We thank Life Science Editors for editing this manuscript. RD and SQ contributed equally to this project and may list themselves first in bibliographic documents. RD, SQ, ZC, and DVV conceived the project. RD and SQ performed algorithm design and implementation for cryosearch. SQ performed data sourcing, filtering, and manual docking/inspections of cryo-ET data. ZC and DVV supervised the project.

## DECLARATIONS

D.V.V. is a scientific cofounder of Aizen Therapeutics and holds equity in the company. All other authors declare no competing interests.

## CODE AVAILABILITY

All code, sample MRC data files, and usage instructions are available at https://github.com/vanvalenlab/cryosearch.

## METHODS

### Implementation and GPU-accelerated docking

Cryosearch was implemented in PyTorch, with all computationally intensive operations— including rasterization, Fast Fourier Transforms (FFTs), and peak extraction—executed on graphics processing units (GPUs). For a given input density map *A*(*x*) and a candidate structure with atomic coordinates 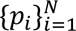, poses defined by a rotation *R* ∈ *SO*(3) and translation *t* ∈ ℝ^3^ were scored by maximizing an FFT-computed cross-correlation over translations for each sampled rotation.

### Target preprocessing, filtering, and model rasterization

To remove background noise prior to docking, the input map was masked using a user-specified density cutoff threshold (*-cutoff*) *c*, generating a thresholded map *A*_0_(*x*):

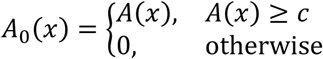

A binary support mask was then defined as *M*(*x*) = I[*A*_0_(*x*) > 0].

To emphasize spatial frequencies consistent with the input resolution, a resolution-aware Fourier-domain filter *F*(*k*) was applied to both the map and the candidate structures. Letting res denote the user-provided resolution (-res, in Å) and *dx* the voxel spacing, the standard deviation of the sampling kernel was approximated as 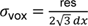. The filter supports two modes:

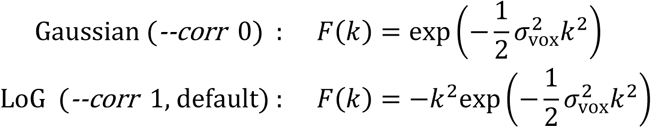

In LoG mode, the zero-frequency component was set to zero (*F*(0) ← 0) to reduce sensitivity to constant offsets and enhance local contrast. The filtered target volume *A*_*f*_(*x*) was computed as:

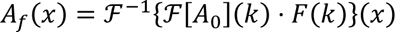

This volume was subsequently normalized to a unit *L*_2_ norm with a small *ϵ* for numerical stability:

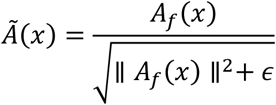

The support mask was reapplied as *Ã*(*x*) = *Ã*(*x*) *M*(*x*). For computational efficiency, the FFT of the processed target was precomputed and cached as 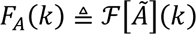 for reuse across all candidate orientations.

### FFT-based docking over translations and rotations

During search, centered atomic coordinates were rotated via Z-Y-Z Euler angles (*q*_*i*_(*R*) = *R p*_*i*_) and rasterized onto the target grid to form a model occupancy volume *B*_*R*_(*x*) using trilinear splatting. Each atom contributed to its eight neighboring voxels weighted by the product of linear barycentric weights. For a fixed rotation *R*, the cross-correlation score over all voxel translations *Δ* was computed using the convolution theorem:

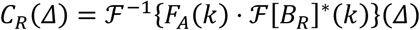

where (⋅)^∗^ denotes complex conjugation. The 3D volume *C*_*R*_(Δ) provided a score for every discrete translation Δ on the grid, obtained with one forward FFT of *B*_*R*_ and one inverse FFT per rotation.

In practice, many rotations in parallel were evaluated using batched FFTs on GPU, and a coarse-to-fine angular strategy was used by default to refine only the highest-scoring coarse orientations.

For each rotation, the best translation was identified as the maximizer of *C*_*R*_ on the discrete grid. Because FFT correlation yields indices in [0, *n* − 1] per axis, each peak index was converted to a signed voxel displacement by wrap-around:

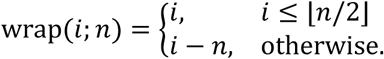

The resulting displacement (in voxels) was converted to physical units using voxel spacing to report *t* in Å. Multiple high-scoring peaks per rotation were retained to produce a ranked list of candidate poses (*R*, *t*) for downstream filtering and multimer assembly.

### Monte Carlo Tree Search (MCTS) assembly and reward function

Multi-component complex assembly was formulated as a Monte Carlo Tree Search (MCTS). A search state *s* consists of a set of frozen component placements 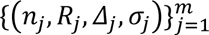, where *σ*_*j*_ ∈ {+1, −1} encodes the chosen translation sign convention. The predicted complex density is given by:

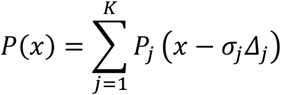

To expand the tree, the algorithm evaluates candidate proteins against the current frozen complex. Let *P*_fixed_ denote the current frozen prediction and *P*_new_(⋅ −*Δ*) denote a candidate probe at translation *Δ*. Candidate evaluation maximizes the normalized correlation *S*(*Δ*) of the combined density with the target map *A*:

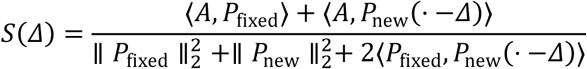

For each candidate addition during assembly, rigid-body docking was performed using a two-stage angular search. A coarse Euler grid (*–deg-coarse*, default 20) was evaluated first, then the top coarse orientations (–top-k, default 100) ranked by docking criteria were subjected to local refinement at 10^∘^angular spacing (*–deg-fine*, default 10). The scalar value backpropagated through the MCTS tree is the terminal reward *R*(*s*), defined as the global normalized cross-correlation:

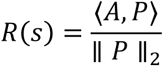

The candidate proteins for MCTS were selected from the top-ranking monomers in the initial screening process, in which monomer docking was performed against all structures in the database and highly correlated monomers were retained as candidates (*–monomer-top*, default 30). Monomer-screen results were cached to avoid repetitive computation during subsequent runs.

The search tree used in Cryosearch was structurally different from the deeper game trees commonly encountered in game applications of MCTS. Tree depth was determined primarily by the number of components to be assembled, whereas tree width was determined by the number of candidate protein models considered at each step. As a result, proteome-scale searches were represented by trees that were relatively shallow but very broad (Extended Data. Fig 8). In large-complex modeling, independent rollouts were therefore often observed to recover different partial assemblies, because early placement decisions strongly constrained the remaining compatible solutions.

### Steric penalization and overlap suppression

To prevent unphysical configurations, soft steric penalties and overlap suppression were applied during docking. For each frozen component, an unfiltered occupancy map was dilated using a spherical top-hat kernel *K*_*r*_ of a defined steric radius (*–steric-radius*, in Å, default 2.0) to establish an excluded-volume region *F*(*x*) = (*O*_fixed_ ∗ *K*_*r*_)(*x*). The candidate’s steric overlap map *C*_steric_(*Δ*) was computed via FFT correlation:

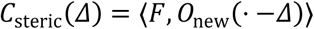

A per-translation clash fraction was defined by normalizing the steric overlap by the candidate’s total occupancy mass. Candidate scores were softly penalized once this fraction exceeded a predefined clash threshold of fractions of atoms in (*–clash-frac*, default 0.2):

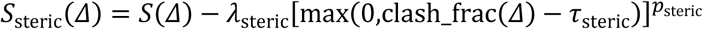

An additional positive overlap term ∝ ⟨*P*_fixed_, *P*_new_(⋅ −*Δ*)⟩ operating on the filtered densities was used to prevent multiple components from claiming the same map features.

### Preparation of the Bos taurus AlphaFold database

The *Bos taurus* reference proteome was downloaded from UniProt (proteome ID UP000009136; taxon ID 9913)^1^. To construct a non-redundant protein library, the UniProt “one protein sequence per gene” set was used as the query database with 22,666 UniProt entries. Corresponding AlphaFold structural models and associated PAE files were then obtained from the AlphaFold Protein Structure Database^2^. All corresponding AlphaFold structures and predicted alignment error (PAE) files were downloaded and processed using *phenix.process_predicted_model*^3,4^. This script removed low-confidence regions (pLDDT ≤ 70) and split the remaining structures into a maximum of five discrete domains per structure (*maximum_domains*=5). The resulting dataset comprised 38,141 domains. Domains shorter than 50 residues were excluded as small domains contain limited structural and shape information that could reduce the specificity in docking. The curation resulted in a final database of 26,136 domains derived from 18,137 gene entries (4,529 gene entries are removed due to parsed domain size under 50 residues). This final database was served as the search database for all docking experiments.

### Docking of the segmented radial spoke map

As a test, the target map was manually cropped based on known trimeric radial spoke head region from the cryo-EM map EMD-50664, resized to a box size of 110 voxels at the original voxel size of 1.3Å to reduce computational cost^5^. The map and models were processed under the resolution reported (*-res* 3.9) and density cutoff of 0.1 (*-cutoff* 0.1) determined in UCSF ChimeraX through visual inspection of the proper level of well-resolved structural features^6^.

An initial exhaustive monomer docking of the map was performed against the *Bos taurus* database, from which the top 30 candidates were retained (*–monomer-top* 30). Multimer assembly was performed via MCTS for 50-500 iterations (*–iters* 50-500) for a target size of three components *(–n components* 3). Steric regularization was enforced by default parameters above. The search retained the top 30 solutions (*–keep-top* 30), and the corresponding complexes were written as output. Modeled assemblies were compared with the reference structure (PDB: 9FQR) to assess whether the expected components were correctly identified and placed within the density map^5^. For convergence analysis, 10 independent MCTS runs were performed. Output models were evaluated based on recovery of the expected trimeric assembly, with complexes containing RSPH1, RSPH9, and either RSPH6A or its close structural homolog RSPH4A scored as successful solutions. If the expected assembly was obtained, the run was defined as successful. The top five outputs from each run were evaluated across 10 runs, and recovery was defined as the percentage of succussful solutions among the 50 total outputs (Extended Data Fig. 3a). Mean wall-clock runtime and standard deviation across the 10 runs for the MCTS on a single NVIDIA RTX A6000 GPU were measured for comparison (Extended Data Fig. 3b).

### COLORES docking

Docking of the segmented radial spoke trimer map (EMD-50664) using the parsed *Bos taurus* domain database was performed using COLORES^5,7,8^, as previously developed for *de novo* protein identification^9^. An angular sampling step of 10° was used to identify the best pose. Runtime for COLORES was measured on a computational cluster equipped with Intel Xeon E5-2687W v3 CPUs. The total run time was estimated as aggregate CPU time across docking jobs in parallel, which was compared to the wall-clock time for the MCTS on a single NVIDIA RTX A6000 GPU (Fig. 1d).

### Preparation of maps of different sizes for monomer docking

Target maps of different sizes were generated by unsupervised segmentation in UCSF ChimeraX Segger using iterative grouping with grouping size of two^6^. One, two, and three rounds of grouping were used to generate maps containing 6, 9, and 12 components, respectively. Each segmented map was then low-pass-filtered to 10 Å, resampled to 2.6 Å/voxel and a box size of 110 voxels to reduce computational cost. Monomer docking was performed using the same docking settings described for the segmented radial spoke map (Extended Data Fig. 1)^5^.

### Docking of the radial spoke 1 head complex

For large-scale assembly modeling, the radial spoke 1 head map was segmented from the composite map (EMD-50664) based on the locally refined EMD-50868 map. This map was then low-pass filtered to 10 Å, resampled to 2.6 Å/voxel and a box size of 110 voxels to reduce computational cost. For monomer docking, the map and models were processed under the resolution parameter of 10.0 Å (*-res* 10.0) and density cutoff of 0.1 (*-cutoff* 0.1) determined in UCSF ChimeraX through visual inspection of the proper level of well-resolved structural features^6^. Following exhaustive monomer docking against the *Bos taurus* database, the top 1,500 candidates were retained for the MCTS assembly (*–monomer-top* 1500). Multimer assembly was optimized over 2,000 iterations (*–iters* 2000) for an estimated 40-component target complex (*–n components* 40). The top 500 solutions were written as output (*–keep-top* 500).

The greedy combination and a 2^nd^ MCTS were employed for further refinement. Models from the top 100 MCTS models were processed through the greedy combination, yielding an intermediate 46-chain complex. The first 40 monomers in the greedy combination were used as the initial MCTS solution for the 2^nd^ MCTS refinement. The 2^nd^ MCTS refinement was run under the same settings as above with the 46 candidate chains as the search database to assemble the final 40-complex model (*–n-components* 40) over 1000 iterations (*–iters* 1000). To evaluate convergence, the greedy combination was performed using increasing numbers of top-ranked MCTS solutions, ranging from 50 to 500 (Extended Data Fig. 6). Modeled assemblies were compared with the reference structure (PDB: 9FQR) to assess recovery of expected protein entries and chains.

### Local resolution analysis

Local resolution was estimated in RELION-5 using the half maps associated with EMD-50868 (raw data of radial spoke 1 head local refinement before sharpening)^10^. The local resolution map was displayed in UCSF ChimeraX^6^.

---

**Extended Data Figure 1.**
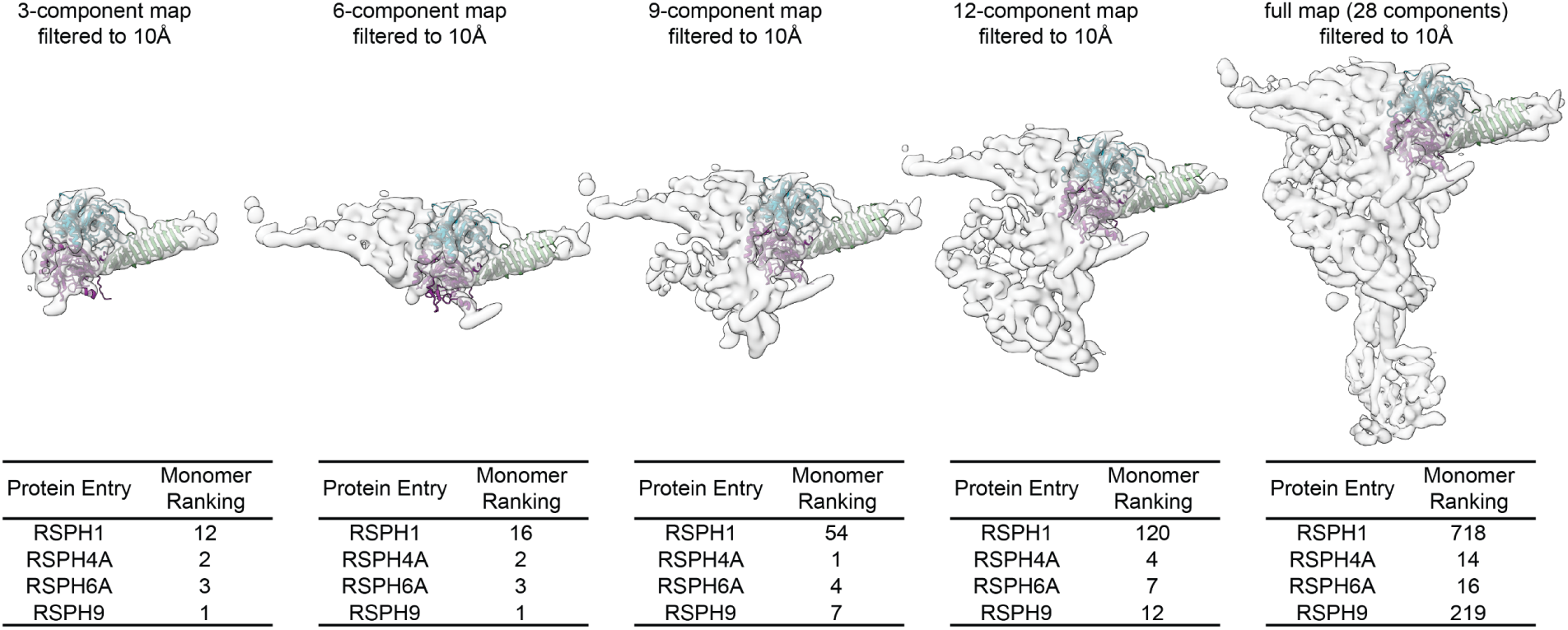
Monomer docking in larger maps yielded more false positives. Comparison of proteome-scale monomer docking rankings for RSPH9, RSPH4A/6A and RSPH1 across maps of increasing volumes (segmented from EMD-50664), showing decreased ranking of the correct components due to false-positives.

**Extended Data Figure 2.**
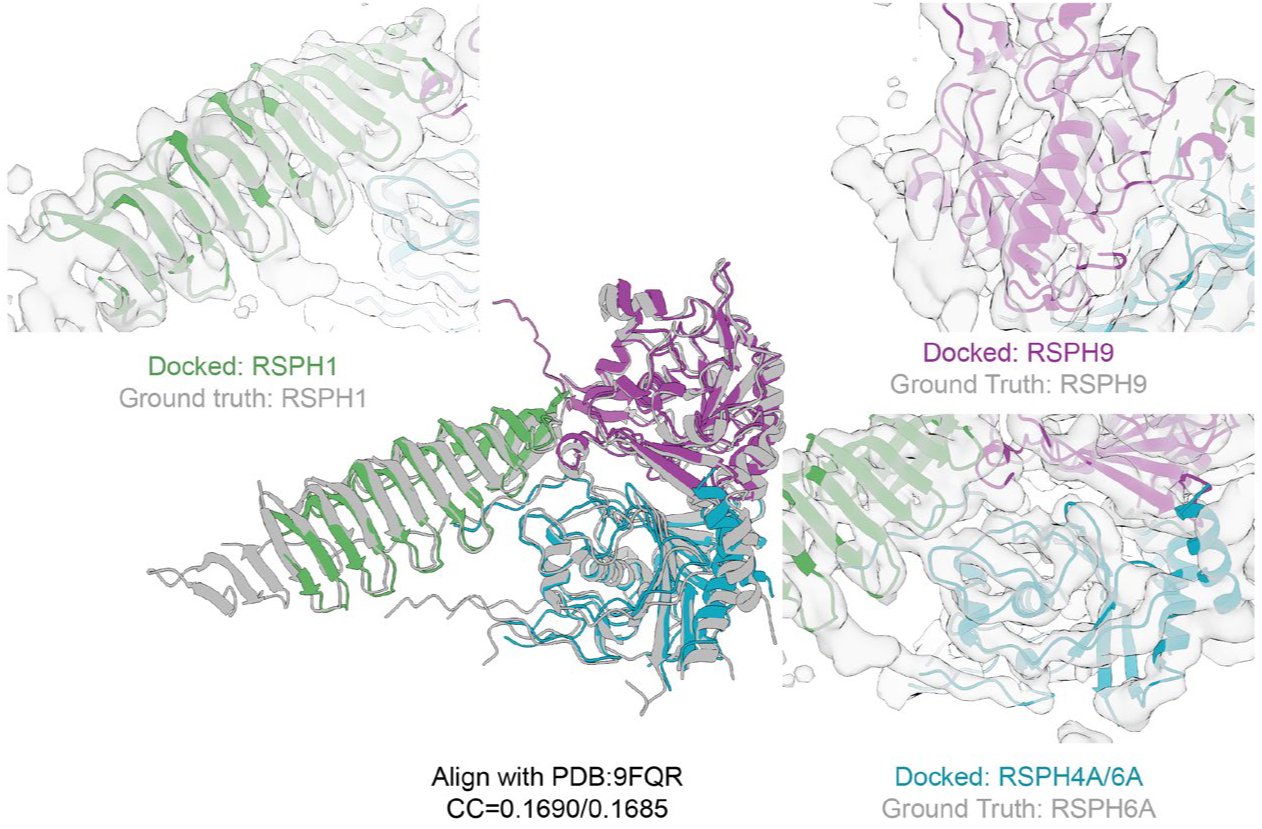
The docked trimer model from Cryosearch is highly consistent with the map features and the reference model. Overlays of the docked trimer models with the reference model (PDB: 9FQR) yielded TM-scores of 0.8629 and 0.8706 (see METHODS). Zoomed-in views of each component fitted into the EMD-50664 map show agreement with secondary-structure features.

**Extended Data Figure 3.**
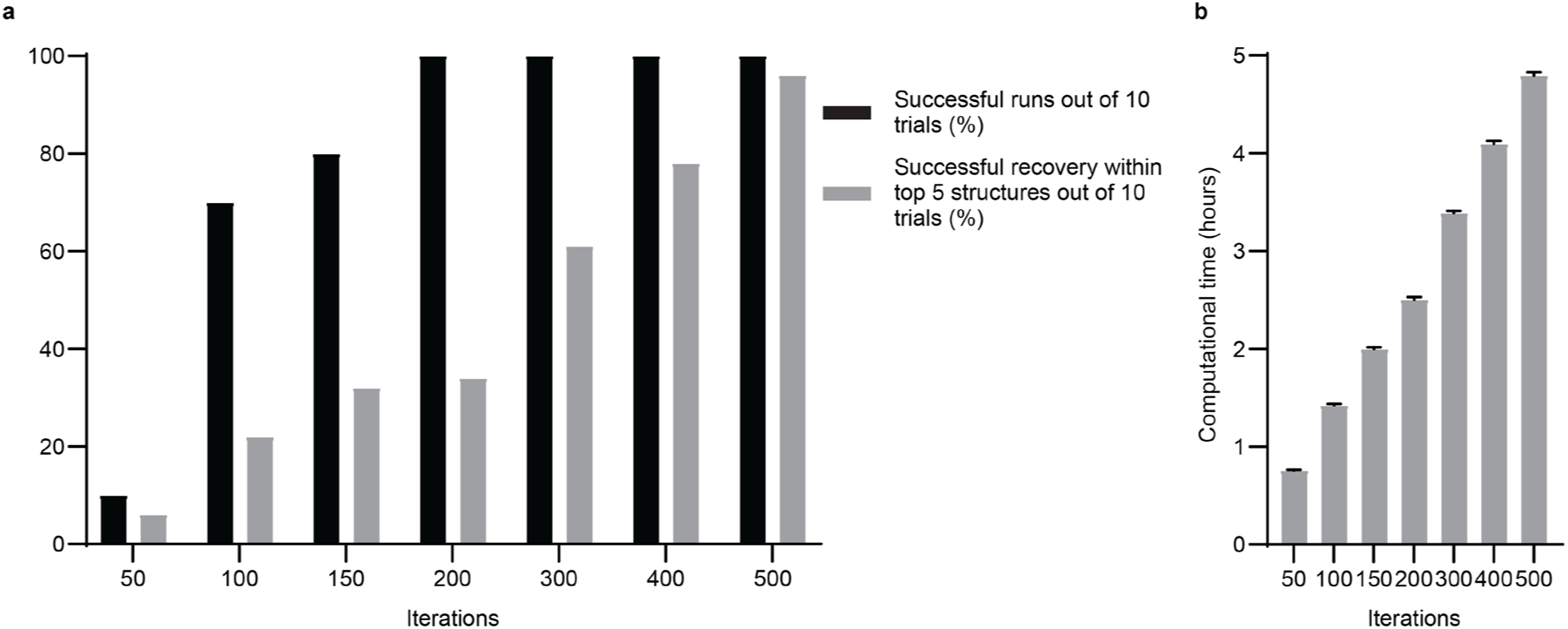
Increasing MCTS iterations improved recovery of the correct assembly with linear runtime scaling. **(a)** Recovery of the correct radial spoke head trimer across 10 independent MCTS runs at varying iteration numbers, showing both run-level success and enrichment of correct assemblies among the top-ranked solutions. **(b)** Runtime at varying iteration numbers on an NVIDIA RTX A6000 GPU, (mean ± s.d. N = 10 independent runs).

**Extended Data Figure 4.**
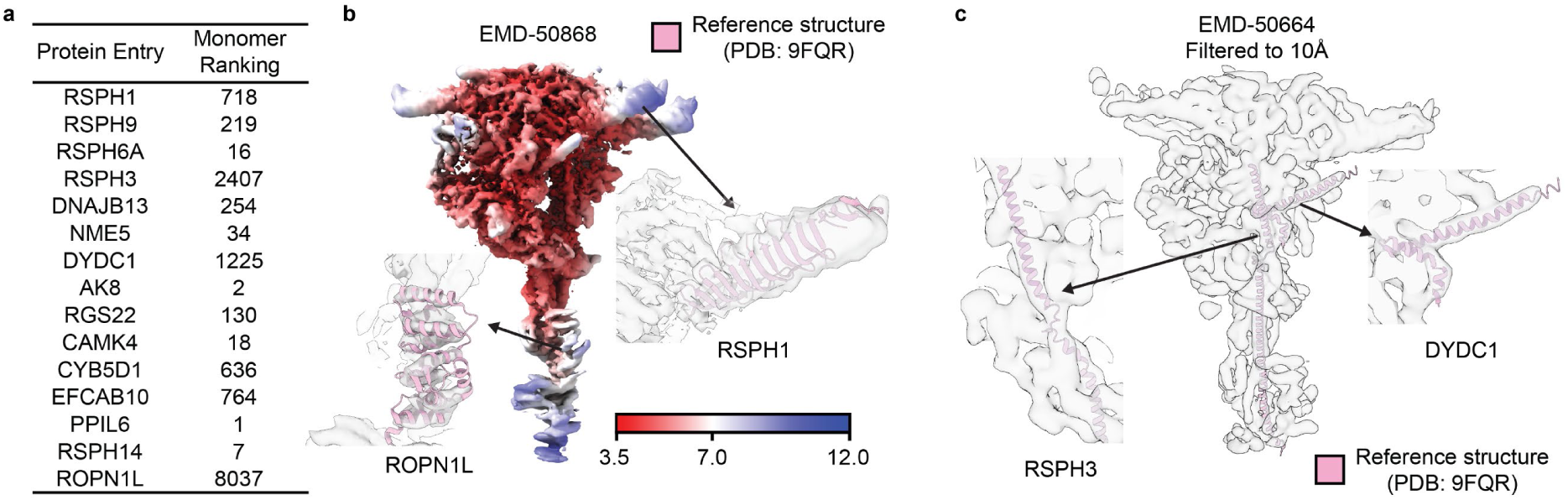
Monomer docking in the low-pass-filtered radial spoke 1 head map, and analyses of low-ranking components due to low local resolution and structural complexity. **(a)** Monomer docking rankings of radial spoke 1 components against the full radial spoke 1 map low-pass filtered to 10 Å (EMD-50664). **(b)** Local resolution estimation of radial spoke 1 (EMD-50868), which was used to build the final composite map (EMD-50664), reveals low-resolution regions corresponding to ROPN1L and RSPH1. **(c)** RSPH3 and DYDC1, comprised of single α-helical features, docked into the map (EMD-50664).

**Extended Data Figure 5.**
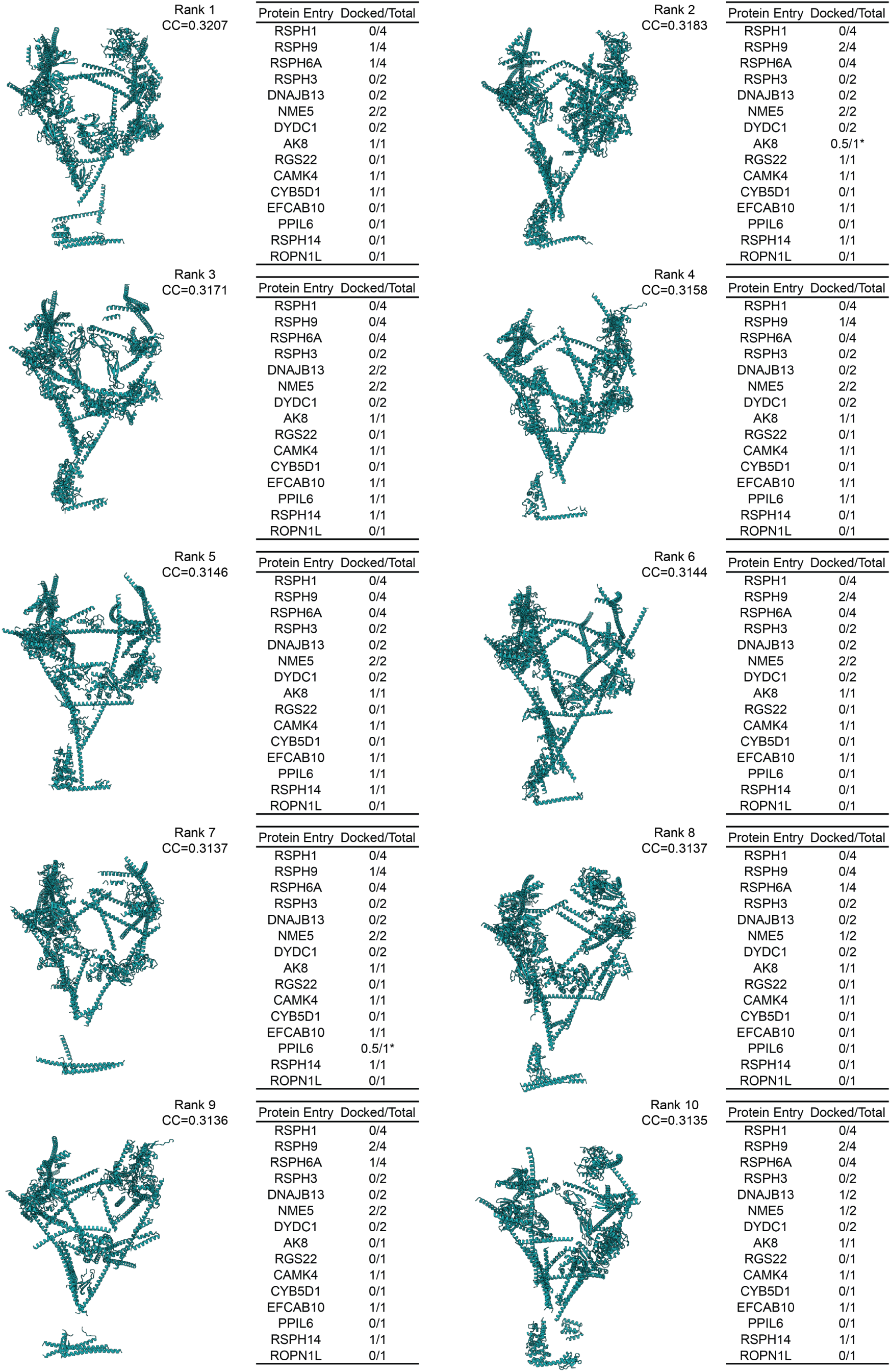
The initial MCTS search recovered partial assemblies. The top 10 complexes and their cross-correlation scores from the initial MCTS search are shown. Different subsets of the correct components were recovered. Asterisks denote that two-domain proteins for which only one domain was recovered.

**Extended Data Figure 6.**
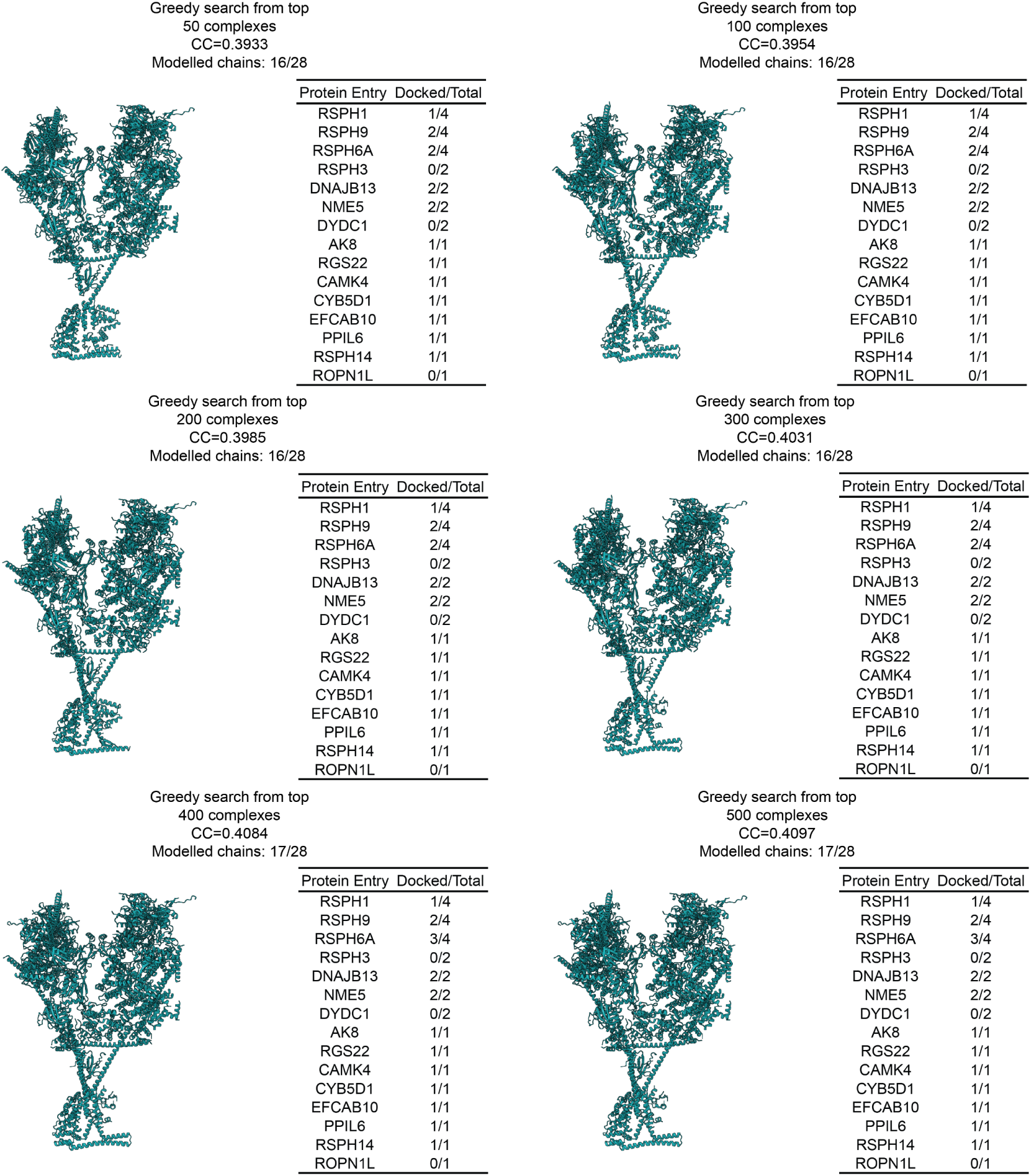
A greedy combination of top-ranked partial assemblies increased the recovery of correct components. Greedy combination results using the top 50, 100, 200, 300, 400 and 500 MCTS solutions, showing recovery of most protein entries (12 out of 15).

**Extended Data Figure 7.**
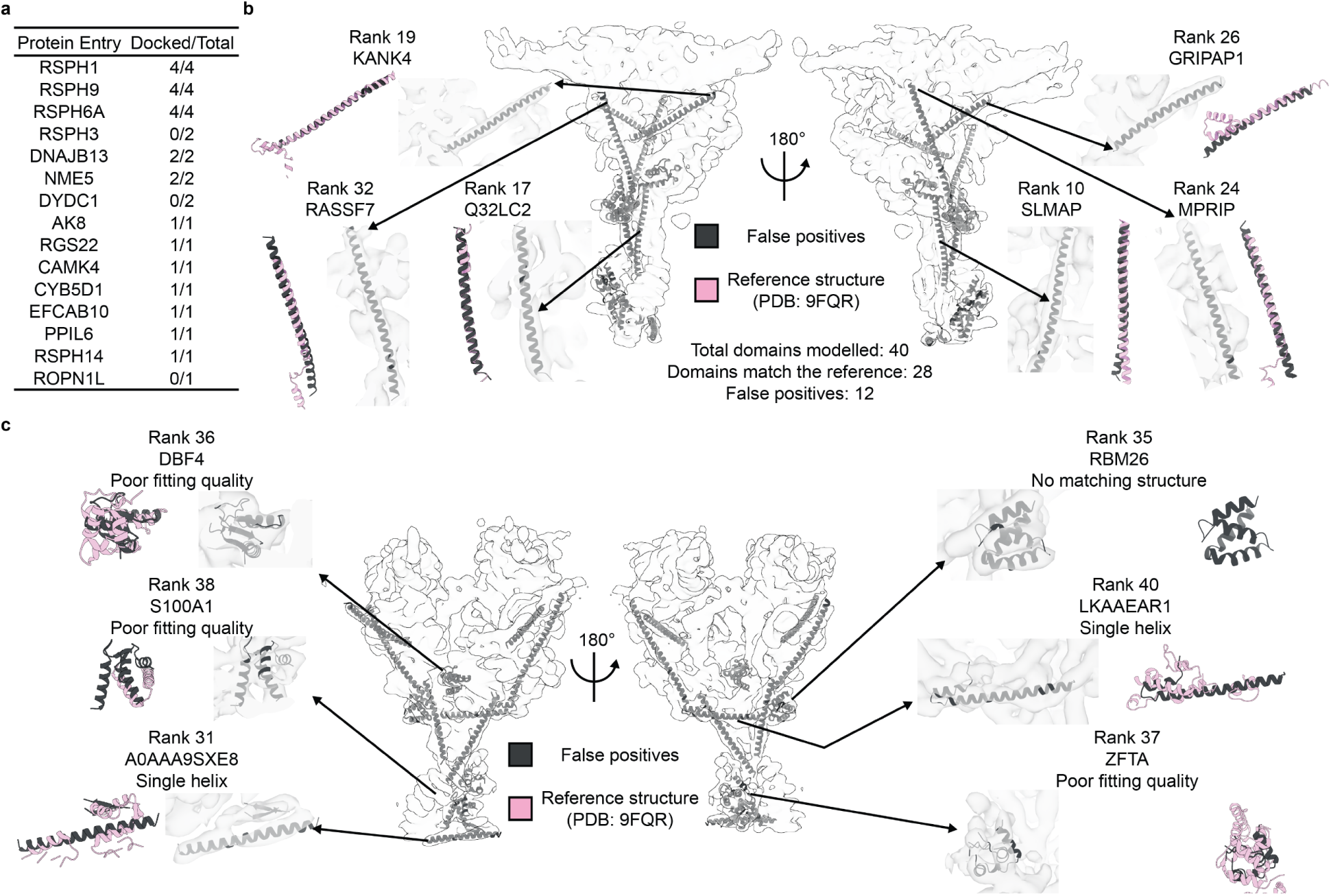
Analyses of false positives. **(a)** Summary of correctly modeled protein entries and chains after the 2^nd^ MCTS refinement. **(b)** Single α-helical models were docked into map regions corresponding to RSPH3 and DYDC1, with their rankings listed. **(c)** Additional false positives in the final assembly highlight a mismatch between map features and candidates.

**Extended Data Figure 8.**
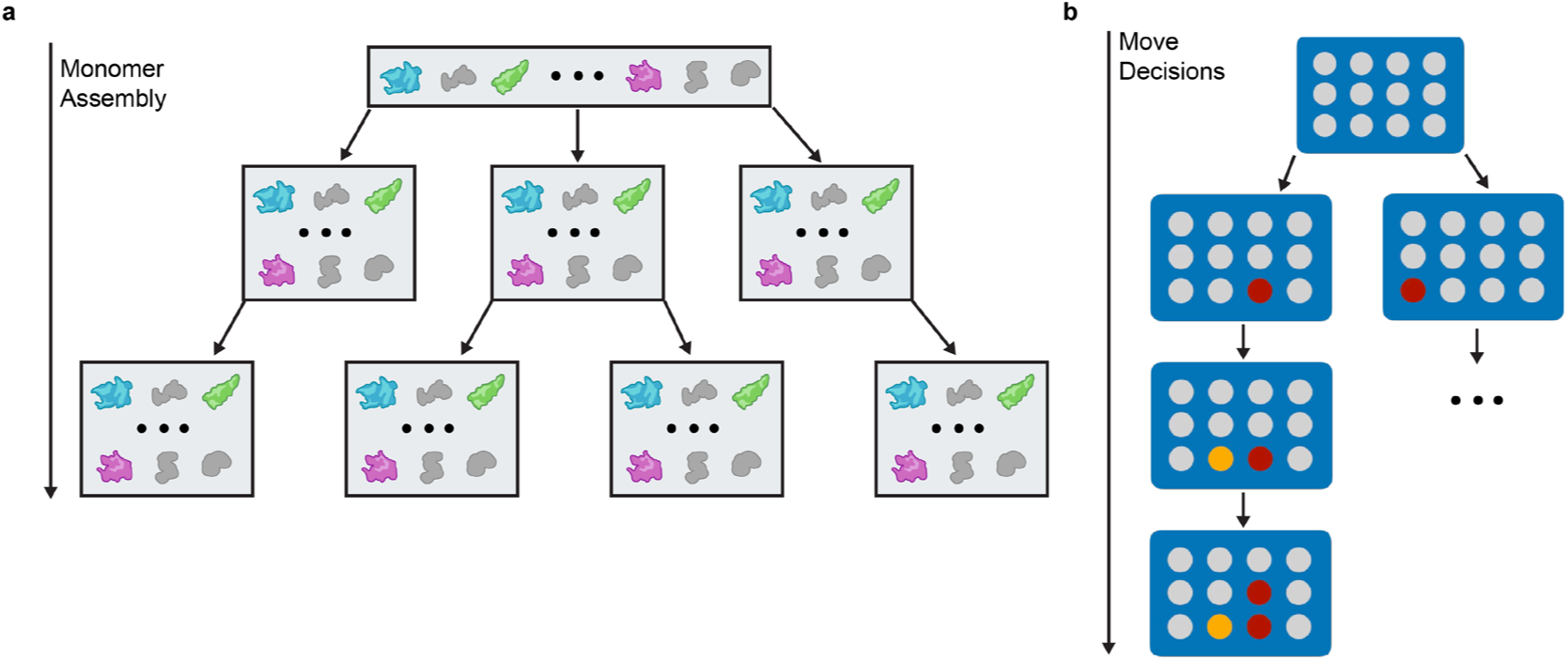
Tree search in complex modeling is shallow and wide compared with trees in games. **(a)** Schematic of protein search profile; the depth is the number of components, and the width of each branch is the size of the monomer database. **(b)** Schematic of game trees, which are deeper but with a reduced number of moves at each branch.

**Extended Data Figure 9.**
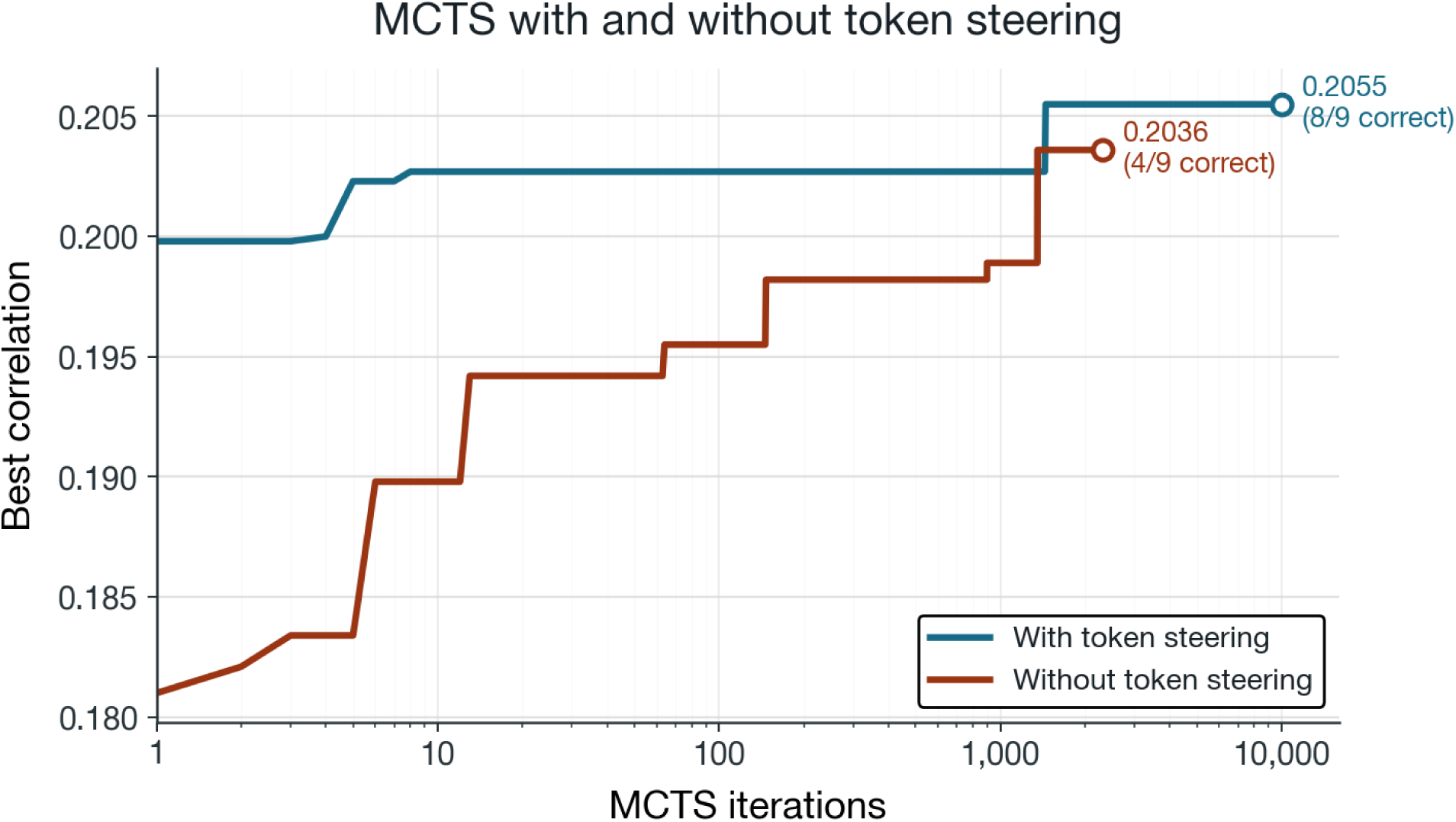
Neural components can accelerate search. By biasing the search algorithm with structural tokenizers, we can converge much more quickly with MCTS. In this example, the complex contains a highly symmetric component, so structural similarity considerably reduces the width of each level in the search tree.

## REFERENCES

1. Nickell, S., Kofler, C., Leis, A. P. & Baumeister, W. A visual approach to proteomics. Nat Rev Mol Cell Biol 7, 225–230 (2006).

2. Bäuerlein, F. J. B. & Baumeister, W. Towards Visual Proteomics at High Resolution. Journal of Molecular Biology 433, 167187 (2021).

3. Cheng, Y. Single-particle cryo-EM—How did it get here and where will it go. Science 361, 876–880 (2018).

4. Briggs, J. A. Structural biology in situ—the potential of subtomogram averaging. Current Opinion in Structural Biology 23, 261–267 (2013).

5. Schur, F. K. Toward high-resolution in situ structural biology with cryo-electron tomography and subtomogram averaging. Current Opinion in Structural Biology 58, 1–9 (2019).

6. Xue, L. et al. Visualizing translation dynamics at atomic detail inside a bacterial cell. Nature 610, 205–211 (2022).

7. Von Kügelgen, A. et al. In Situ Structure of an Intact Lipopolysaccharide-Bound Bacterial Surface Layer. Cell 180, 348–358.e15 (2020).

8. Watson, A. J. I. & Bartesaghi, A. Advances in cryo-ET data processing: meeting the demands of visual proteomics. Current Opinion in Structural Biology 87, 102861 (2024).

9. Casañal, A., Shakeel, S. & Passmore, L. A. Interpretation of medium resolution cryoEM maps of multi-protein complexes. Current Opinion in Structural Biology 58, 166–174 (2019).

10. Chen, Z. et al. De novo protein identification in mammalian sperm using in situ cryoelectron tomography and AlphaFold2 docking. Cell 186, 5041–5053.e19 (2023).

11. Gao, J. et al. DomainFit: Identification of protein domains in cryo-EM maps at intermediate resolution using AlphaFold2-predicted models. Structure 32, 1248–1259.e5 (2024).

12. Ma, M. et al. Structure of the Decorated Ciliary Doublet Microtubule. Cell 179, 909–922.e12 (2019).

13. Gui, M. et al. Structures of radial spokes and associated complexes important for ciliary motility. Nat Struct Mol Biol 28, 29–37 (2021).

14. Jamali, K. et al. Automated model building and protein identification in cryo-EM maps. Nature 628, 450–457 (2024).

15. Terwilliger, T. C., Adams, P. D., Afonine, P. V. & Sobolev, O. V. Cryo-EM map interpretation and protein model-building using iterative map segmentation. Protein Science 29, 87–99 (2020).

16. Jumper, J. et al. Highly accurate protein structure prediction with AlphaFold. Nature 596, 583–589 (2021).

17. Chacón, P. & Wriggers, W. Multi-resolution contour-based fitting of macromolecular structures. Journal of Molecular Biology 317, 375–384 (2002).

18. Leung, M. R. et al. Structural diversity of axonemes across mammalian motile cilia. Nature 637, 1170–1177 (2025).

19. Varadi, M. et al. AlphaFold Protein Structure Database: massively expanding the structural coverage of protein-sequence space with high-accuracy models. Nucleic Acids Research 50, D439–D444 (2022).

20. Wriggers, W. Conventions and workflows for using *Situs*. Acta Crystallogr D Biol Crystallogr 68, 344–351 (2012).

21. Bryant, P. et al. Predicting the structure of large protein complexes using AlphaFold and Monte Carlo tree search. Nat Commun 13, 6028 (2022).

22. Maddhuri Venkata Subramaniya, S. R., Terashi, G. & Kihara, D. Protein secondary structure detection in intermediate-resolution cryo-EM maps using deep learning. Nat Methods 16, 911–917 (2019).

23. Kucukelbir, A., Sigworth, F. J. & Tagare, H. D. Quantifying the local resolution of cryo-EM density maps. Nat Methods 11, 63–65 (2014).

24. Silver, D. et al. Mastering the game of Go with deep neural networks and tree search. Nature 529, 484–489 (2016).

25. Dilip, Rohit, Ayush Varshney, and David Van Valen. "Adaptive Protein Tokenization." arXiv preprint arXiv:2602.06418 (2026).

## REFERENCES

1. The UniProt Consortium et al. UniProt: the Universal Protein Knowledgebase in 2025. Nucleic Acids Research 53, D609–D617 (2025).

2. Varadi, M. et al. AlphaFold Protein Structure Database: massively expanding the structural coverage of protein-sequence space with high-accuracy models. Nucleic Acids Research 50, D439–D444 (2022).

3. Oeffner, R. D. et al. Putting *AlphaFold* models to work with phenix.process_predicted_model and ISOLDE. Acta Crystallogr D Struct Biol 78, 1303–1314 (2022).

4. Liebschner, D. et al. Macromolecular structure determination using X-rays, neutrons and electrons: recent developments in *Phenix*. Acta Crystallogr D Struct Biol 75, 861–877 (2019).

5. Leung, M. R. et al. Structural diversity of axonemes across mammalian motile cilia. Nature 637, 1170–1177 (2025).

6. Meng, E. C. et al. UCSF CHIMERAX : Tools for structure building and analysis. Protein Science 32, e4792 (2023).

7. Chacón, P. & Wriggers, W. Multi-resolution contour-based fitting of macromolecular structures. Journal of Molecular Biology 317, 375–384 (2002).

8. Wriggers, W. Conventions and workflows for using *Situs*. Acta Crystallogr D Biol Crystallogr 68, 344–351 (2012).

9. Chen, Z. et al. De novo protein identification in mammalian sperm using in situ cryoelectron tomography and AlphaFold2 docking. Cell 186, 5041–5053.e19 (2023).

10. Burt, A. et al. An image processing pipeline for electron cryo-tomography in RELION-5. FEBS Open Bio 14, 1788–1804 (2024).

